# Consensus Phylogenetic trees of Fifteen Prokaryotic Aminoacyl-tRNA Synthetase Polypeptides based on Euclidean Geometry of All-Pairs Distances and Concatenation

**DOI:** 10.1101/051623

**Authors:** Rhishikesh Bargaje, M.Milner Kumar, Sohan Prabhakar Modak

## Abstract

**Background:** Most molecular phylogenetic trees depict the relative closeness or the extent of similarity among a set of taxa based on comparison of sequences of homologous genes or proteins. Since the tree topology for individual monogenic traits varies among the same set of organisms and does not overlap taxonomic hierarchy, hence there is a need to generate multidimensional phylogenetic trees.

**Results:** Phylogenetic trees were constructed for 119 prokaryotes representing 2 phyla under Archaea and 11 phyla under Bacteria after comparing multiple sequence alignments for 15 different aminoacyl-tRNA synthetase polypeptides. The topology of Neighbor Joining (NJ) trees for individual tRNA synthetase polypeptides varied substantially. We use Euclidean geometry to estimate all-pairs distances in order to construct phylogenetic trees. Further, we used a novel “Taxonomic fidelity” algorithm to estimate clade by clade similarity between the phylogenetic tree and the taxonomic tree. We find that, as compared to trees for individual tRNA synthetase polypeptides and rDNA sequences, the topology of our Euclidean tree and that for aligned and concatenated sequences of 15 proteins are closer to the taxonomic trees and offer the best consensus. We have also aligned sequences after concatenation, and find that by changing the order of sequence joining prior to alignment, the tree topologies vary. In contrast, changing the types of polypeptides in the grouping for Euclidean trees does not affect the tree topologies.

**Conclusions:** We show that a consensus phylogenetic tree of 15 polypeptides from 14 aminoacyl-tRNA synthetases for 119 prokaryotes using Euclidean geometry exhibits better taxonomic fidelity than trees for individual tRNA synthetase polypeptides as well as 16S rDNA. We have also examined Euclidean N-dimensional trees for 15 tRNA synthetase polypeptides which give the same topology as that constructed after amalgamating 3-dimensional Euclidean trees for groups of 3 polypeptides. Euclidean N-dimensional trees offer a reliable future to multi-genic molecular phylogenetics.

## Introduction

In order to establish evolutionary relationships among a set of taxa, phylogenetic trees are constructed by assessing the extent of similarity among morphological, physiological, biochemical, genetic, and behavioral traits [1] and such trees closely resemble the hierarchy of taxonomic classification. These phylogenetic trees can be further supplemented by comparing the sequence data on nucleic acid (nucleotides) and protein (amino acid) from different organisms. Indeed, rRNA phylogenetic trees have also been used to supplement classical taxonomic trees (NCBI). In contrast, paucity of information on nucleotide sequences of individual genes, open reading frames or mRNAs, as well as amino acid sequences of polypeptides has been responsible for lack of their incorporation into taxonomic trees. Nonetheless, available data is used to construct trees to assess the relative phylogenetic position of taxa from groups with potential evolutionary links. Unfortunately, topology of individual trees for different molecular traits varies considerably [2,3]. In contrast, morphological, genetic, physiological or biochemical traits reflect a combination of multi-genic traits [3]. Indeed, a comparison of entire genome sequences should yield meaningful insight in the evolutionary relationships. Recently, visualization, analysis and comparison of fractal structure of genomes has revealed significant phylogenetic similarities among closely related taxa and differences among distant ones, [4–6] but no attempt has yet been made to interpret positional differences in the fractal diagrams in relation to the discrete portion of genomes.

With the increasing availability of nucleotide-and amino acid sequences, molecular phylogenetic trees for the same set of taxa have been constructed. Yet since these often do not exhibit the same topology as the taxonomic trees, the differences lead to a controversy between classical taxonomy [7] and phylogenetic cladistics [8]. Indeed, evolution manifests in the form of changes in the whole organism and its genetic informational reservoir, not in individual genes, which are subject to random mutations at variable rates and cannot alone affect the principal phenotype of the organism [7]. Instead, one would expect that a number of gene cohorts operating in a concerted manner is what leads to changes in the phenotype of a polygenic trait [3].

Recently, sequences of multiple genes or polypeptides have been concatenated (end-to-end joined) in tandem in order to generate large multi-sequence strings that are used to construct phylogenetic trees [2]. However, this method requires alignment of individual sequences before concatenation. Milner et. al. [3, 9,10] have developed another method that establishes coordinates for the extent of similarity among 3 individual traits or parameters for a set of species. The method positions these coordinates along x,y,z axes to generate consensus all pairs distances to construct and visualize the resultant tree structure in a 3-dimensional space. Recently Milner et. al. used this method to construct a single 3D phylogenetic tree for 3 mitochondrial polypeptides for a set of 76 species and found that the 3D phylogenetic tree closely mimicked the consensus taxonomic tree [11]. It is, therefore, necessary to extend this method to larger number of molecular traits in order to achieve a consensus phylogenetic representation.

Here, using Euclidean geometry [9,10], we have constructed stepwise multiparametric trees in 3D for 15 aminoacyl-tRNA synthetase polypeptides from 119 species from prokaryotes and archaea and compared the tree topology to that of trees for rDNA, individual tRNA synthetases and their concatenates. As our method handles only 3 parameters/polypeptides at a time to generate a Euclidean distance matrix, we grouped tRNA synthetases in groups of 3 according to their chemical and physical properties and then amalgamated these matrices stepwise to obtain a consensus representation for 15 aminoacyl-tRNA synthetases. We have also used a novel N-dimensional metric to obtain all pairs distances for these 15 polypeptides simultaneously to construct a consensus phylogenetic tree and found that it yields topology identical to that using amalgamated 3-polypeptide matrices. Recently, Milner Kumar & Sohan P. Modak (manuscript in review) have compared phylogenetic trees for single polypeptides and multiple polypeptide-trees based on Euclidean geometry to the classical taxonomic trees to assess the extent of fidelity (‘F’ value) or topological similarities for the given set of species. We find that, unlike single polypeptide trees, Euclidean phylogenetic trees based on groups of 3 polypeptides/parameters or 15 polypeptides/parameters exhibit highest F value of 90.3%, as compared to 88.9-89.6% for trees using end-to-end joined sequences, and 87.2% for rDNA tree.

## Results

We selected sequence data for 15 polypeptides representing 14 aminoacyl-tRNA synthetases from 119 species. Note that the numbers of species representing different phyla are not similar. Markedly, out of ten phyla from 2 super kingdoms, Phylum Firmicutes and Proteobacteria are represented by 33 and 51 species, respectively.

## Phylogenetic trees for individual aminoacyl-tRNA synthetase polypeptides

The topology of phylogenetic tree for individual tRNA synthetase was compared to the NCBI benchmark taxonomic tree and their taxonomic fidelity ‘F’ was estimated (see methods and discussed in detail in subsequent section). Fig. 1 shows three representative trees exhibiting highest (Leu), lowest (Val) and intermediate (Thr) ‘F’ values (Table 1).

**Figure 1:**
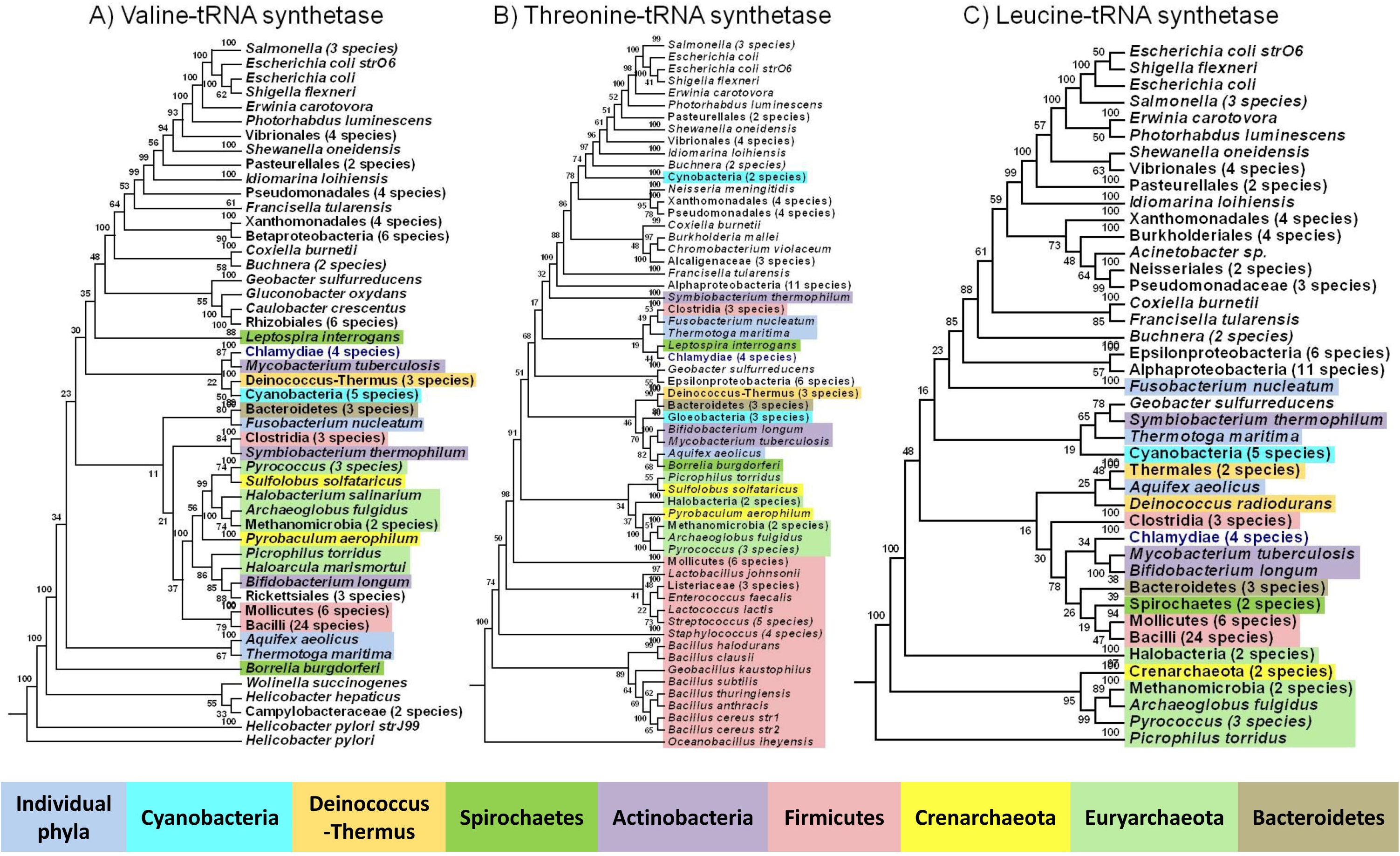
Phylogenetic trees for aminoacyl-tRNA synthetases. Three representative phylogenetic trees with lowest, intermediate and highest ‘F’ value (explained later) are presented in this figure. Note the topological differences among the phylogenetic trees for three aminoacyl-tRNA synthetase. If all species in a particular taxonomic definition form a single clade, such clades are compressed and number of species is mentioned in parenthesis. Expanded version of the trees is available as Supplementary Figure 1. In the Consensus tree the number on each branch indicates the frequency of occurrence from 100 bootstraps. The color code is maintained in remaining figures.

**Table 1:** ‘F’ and ‘RF’ value for phylogenetic trees constructed using various methods.

| Tree | RF value | F Value | Tree | RF value | F Value |
| --- | --- | --- | --- | --- | --- |
| Arg | 113 | 84% | Val | 109 | 83.5% |
| Asp | 103 | 87.3% | Increasing concat-MSA | 99 | 88.9% |
| Cys | 103 | 84% | Decreasing concat-MSA | 99 | 89.6% |
| Ile | 109 | 89.4% | Alphabetical concat-MSA | 103 | 89.2% |
| Leu | 101 | 92.9% | MSA-concat | 99 | 88.9% |
| Lys | 101 | 90.4% | Final Euclidean | 107 | 90.3% |
| Met | 109 | 87.2% | Mean 'D' Euclidean | 107 | 90.3% |
| Phe-α | 101 | 89.7% | Sum 'D' Euclidean | 107 | 90.3% |
| **Phe-β** | 105 | 88% | **Final Euclidean w/o Bootstrap** | 106 | 90.3% |
| **Pro** | 111 | 86.7% | **Mean ‘D’ Euclidean w/o Bootstrap** | 106 | 90.3% |
| **Ser** | 103 | 89.8% | **Sum ‘D’ Euclidean w/o Bootstrap** | 106 | 90.3% |
| **Thr** | 109 | 87.3% | **15 Dimensional** | 107 | 90.3% |
| **Trp** | 111 | 85.1% | **16S rDNA** | 115 | 87.2% |
| **Tyr** | 107 | 85.7% |  |  |  |

Two species from the phylum Crenarchaeota and 9 from the phylum Euryarchaeota (superkingdom Archaea) form unique clade in trees for Asp, Thr and Tyr-aminoacyl-tRNA synthetases. In the remaining 12 trees Archaea were dispersed among Bacteria (Fig. 1 and S1 Figure). Species from the phylum Crenarchaeota form a discrete clade in only six (Arg, Leu. Gly, Ser, Met and Phe-α) trees. In contrast, species from the phylum Euryarchaeota are interspersed with Crenarchaeota. Furthermore, *Borelia* (phylum Spirochaetes), *Bacteroides* and *Chlorobium* (phylum Bacteroidetes) and 3 species from phylum Deinococcus-thermus are intermingled with those from phylum Euryarchaeota (Fig. 1 and S1 Figure). Excepting *Methanosarcina sps*, species belonging to every other genus examined in Archaea form a single clade; *Picrophilus torridus* and *Halobacteria sps*, the lone representatives of their genus, were dispersed among bacterial clades in most of the single gene trees (S1 Figure).

From the superkingdom Bacteria, all 4 species from the phylum Chlamydiae form a discrete clade in all single gene trees. Similarly, 3 species from the phylum Deinococcus-Thermus, 5 from Cyanobacteria and 3 from Bacteroidetes form discrete clades in most trees. In contrast, 2 Spirochaetes are often interspersed with bacteria (Fig. 1 S1 FigureS1 Figure). Finally, the representatives from the phyla Firmicutes, Actinobacteria and Proteobacteria species do not form an independent clade but are dispersed in all trees.

Under the phylum Firmicutes, 3 species from the class Clostridia form a clade, distinct from the clade for the remaining 30 species belonging to the class Mollicutes and Bacilli. In contrast, 45 to 50 species from the phylum Proteobacteria form one or two distinct clades while the rest are interspersed throughout the tree. Conversely, in some trees, species from other phyla are interspersed with Proteobacteria. Taxa from classes Proteobacteria, Epsilonproteobacteria form a single clade in 13 out of 15 tRNA synthetase trees. Classes Alphaproteobacteria and Betaproteobacteria form single clade in almost half the trees while 27 species under the class Gammaproteobacteria never form a distinct clade and are dispersed throughout the trees (Fig. 1 and S1 Figure).

In all trees, the genus *Ureaplasma* fell in the *Mycoplasma* cluster (S1 Figure). Likewise, 3 species under the genus *Chlamydia* joined the clade with *Chlamydophila caviae* (S1 Figure). Finally, in different single polypeptide trees, 7 species from the genus Bacillus form clades in different combinations. We also note that the species under the Genera *Clostridium*, *Escherichia*, *Haemophilus*, *Helicobacter*, *Mycoplasma*, *Pseudomonas* and *Xylella* exhibit variable positions and do not necessarily form individual clades (Fig. 1 S1 Figure).

## Multiparametric phylogenetic trees

### Concatenated tRNA synthetases

We constructed two different types of phylogenetic trees of concatenated sequences. In the first, 15 sequences from 119 species were concatenated to form megapolypeptides and then subjected to multiple sequence alignment (MSA). In the second, each polypeptide for 119 species was first subjected to MSA and then the aligned sequences from 15 polypeptides were concatenated.

All phylogenetic trees generated using the four different concatenation strategies (see methods) show distinct topological positions for the cluster superkingdom Archaea, and the cluster phylum Proteobacteria. The remaining 11 phyla occupy an intermediate position. Five out of ten phyla with at least two representative species (Chlamydiae, Bacteroidetes, Cyanobacteria, Deinococcus-Thermus and Crenarchaeota) form independent clades (Fig. 2; S2 Figure). Under the phylum Firmicutes, species from classes Bacilli, Moillicutes and Clostridia form separate clusters and *Fusobacterium nucleatum* (phylum Fusobacteria) form a clade with Clostridium in all four types of concatenated trees (Fig. 2; S2 Figure). Similarly, species from classes Alphaproteobacteria and Epsilonproteobacteria (phylum Proteobacteria) form single clades. In contrast, Betaproteobacteria form a distinct clade only when sequences are concatenated in either descending or alphabetical orders (concat **→**MSA), but not in the remaining two methods (Fig. 2; S2 Figure).

With all four methods of concatenation, the topology of the top half of the phylogenetic tree remains similar and includes all archaea and bacteria excluding beta-and gamma-proteobacteria (Fig. 2; S2 Figure). The taxa occupying the position near the tree base are Neisseria and Chromobacterium for increasing order concat-MSA and MSA-concat. Xylella appears at the tree base in the alphabetical order concat-MSA, and Salmonella in the decreasing order concat-MSA.

**Figure 2:**
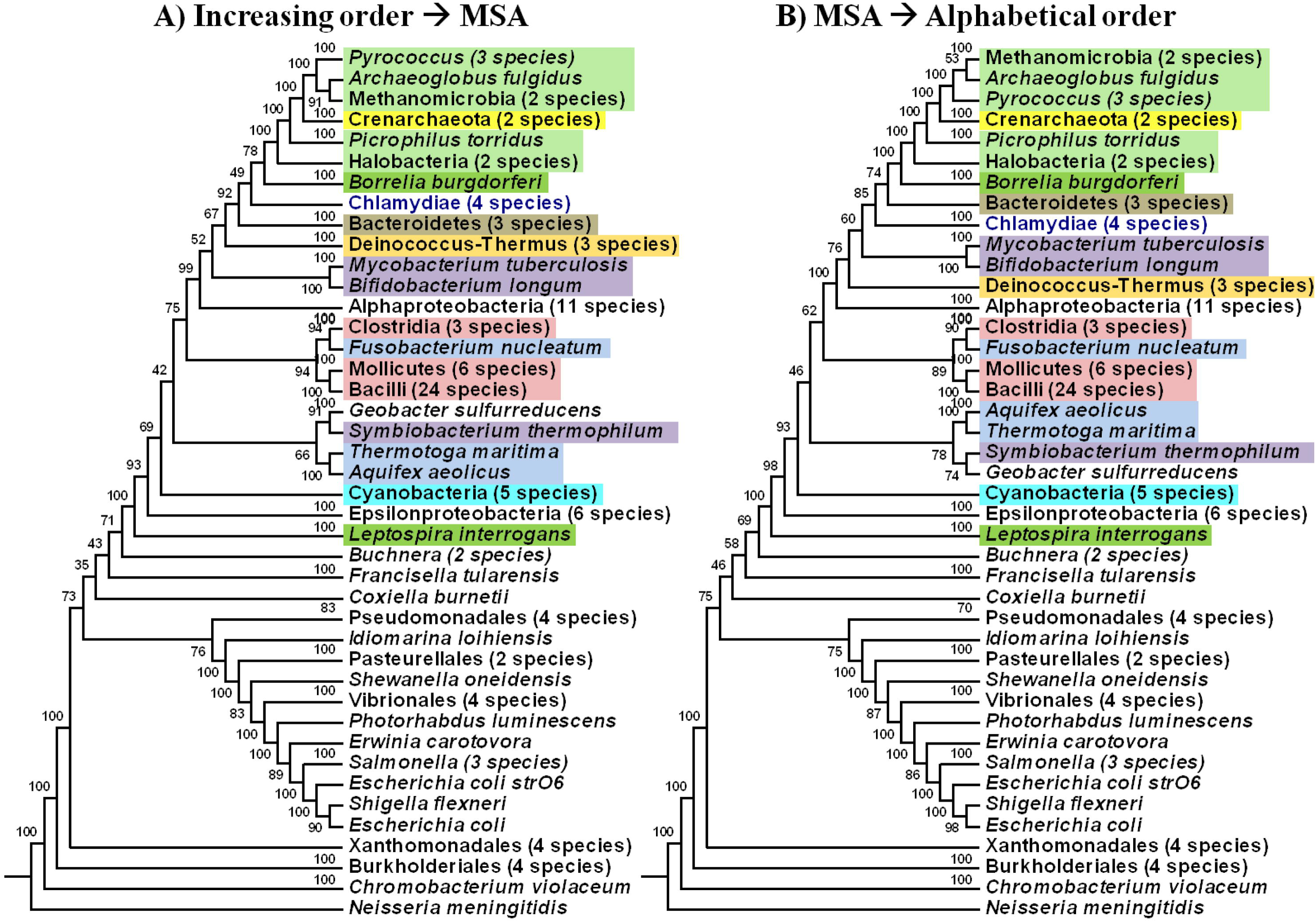
Multidimensional trees using concatenation. Phylogenetic trees constructed using concatenated sequences. The sequences were concatenated in different orders as mentioned in methods section. Expanded version of trees is provided in Supplementary Figure 2.

### Phylogenetic trees for Block (Conserved) and Non-block (Non-conserved) regions of tRNA synthetase sequences

We extracted regions containing conserved and non-conserved sequences of the megapolypeptides using program “GBlocks” [12]. The tree topology of Non-block-concatenated sequences is similar to that of total concatenates (S3 Figure), whereas trees constructed using Block-concatenates showed considerable variation. Topological position of classes Alpha-, Epsilon-Preoteobacteria, Bacilli and Mollicutes varied greatly among trees based on different types concatenation protocols (S3 Figure). In Block-concatenate trees species from classes Gamma-and Beta-proteobacteria are located at the base but with varying topology (S3 Figure). Thus, trees for concatenated Non-block sequences resemble those for total sequences rather than Blocks.

### Phylogenetic trees using Euclidean distances

To construct multiparametric trees in 3D space, 15 aminoacyl-tRNA synthetase polypeptides were first segregated into 5 groups with three molecules each based on three different parameters. These groupings were based on 1) the chemical properties of the acceptor amino acids; 2) the increasing order of sum of all pairs distance for each aminoacyl-tRNA synthetase; and 3) the estimated mean of highest all pairs distance for each species for individual aminoacyl-tRNA synthetase. The proteins were grouped according to their increasing order (see methods). Each method yielded a consensus of 5 trees for 3 proteins, which were then merged stepwise into a single tree (see methods). Notably, all grouping strategies resulted in identical consensus tree topologies (Fig. 3; S4 Figure). We find that the topology of bootstrapped Euclidean distance trees resembles those using concatenation (Fig. 2 and 3).

**Figure 3:**
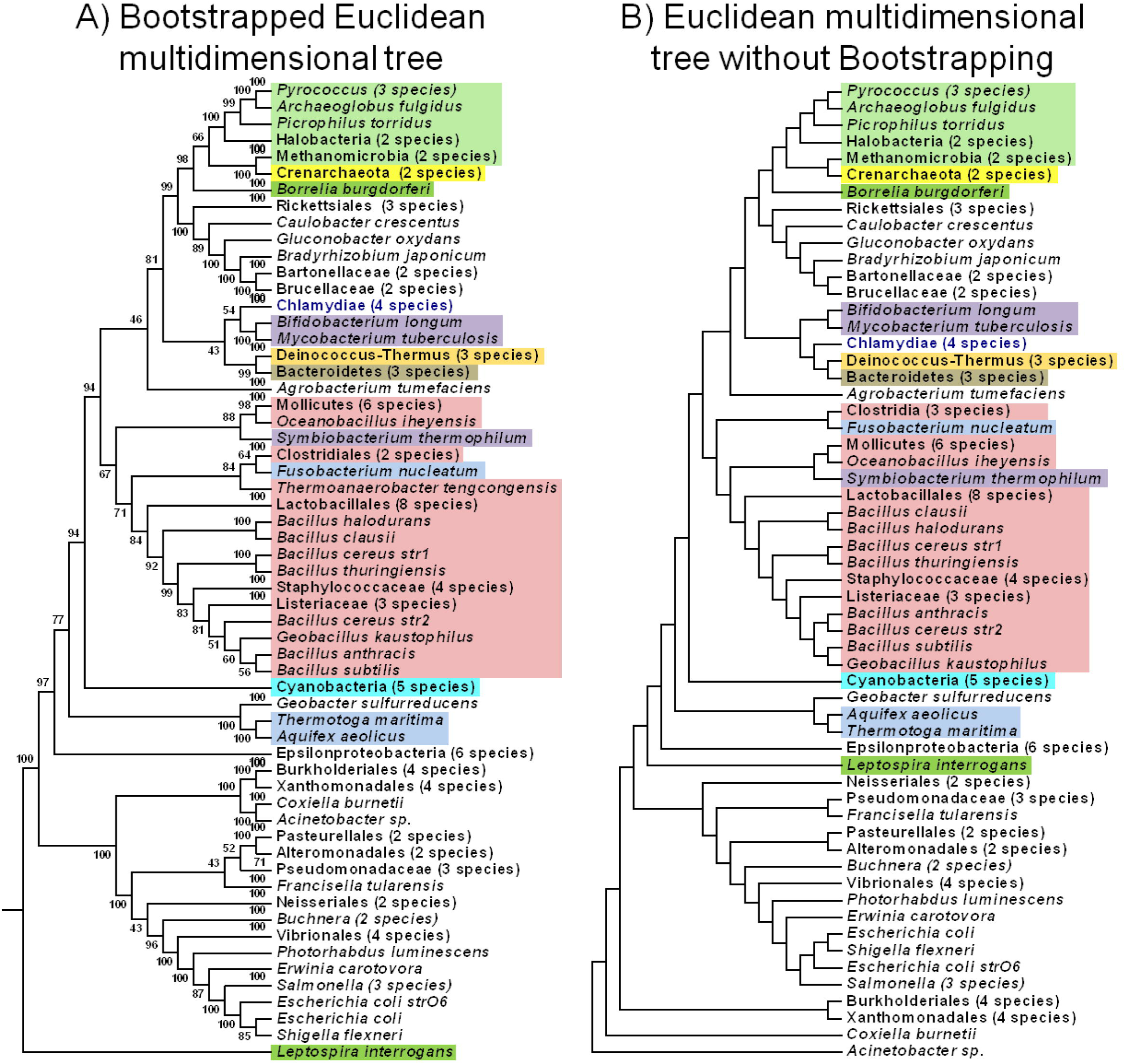
Multidimensional phylogenetic trees for Euclidean distances with or without bootstrapping. Euclidean trees for all three grouping methods (Fig. 6) and 15-dimension resulted in identical trees with the use of bootstrapping (A) and without the use of bootstrapping (B). The expended version of trees is provided as Supplementary Figure 4.

In the Euclidean consensus tree, superkingdom Archaea form a discrete cluster in which species from the phylum Crenarchaeota form a clade with species from class Methanomicrobia of the phylum Euryarchaeota (Fig. 3; S4 FigureS4 Figure). Among 11 bacterial phyla, 52 species under the phylum Proteobacteria are segregated into two major clusters separated by remaining bacterial phyla. We note that species under the phyla Firmicutes, Spirochaetes and Actinobacteria, are dispersed. At the level of phyla the topology of Euclidean trees is considerably similar to those using concatenated sequences, except for differences in the position of species within the clades formed by classes Bacilli, Clostridia, Gloeobacteria, Betaproteobacteria, Alphaproteobacteria and Gammaproteobacteria (Fig. 2 & 3; S2 & S4 Figure). While the species in the phylum Firmicutes form a cluster, *Fusobacterium nucleatum* (Phyluym Fusobacterium) and *Symbiobacterium thermophilum* (phylum Actinobacteria) are interspersed within this group (Fig. 3; S4 Figure). In the phylum Proteobacteria, only Epsilonproteobacteria form a discrete clade. Both species in the family Neisseriaceae (*Neisseria meningitidis* and *Chromobacterium violaceum*) and those from the order Alteromonadales form distinct clades in Euclidean multidimensional trees (Fig. 3; S4 Figure) and resembled the case in the single gene tree for Tyr-tRNA synthetase.

In order to assess if bootstrapping affects the topology of Euclidean trees, we compared the Euclidean trees with and without bootstrapping. We find that there is almost no difference between the topologies of trees constructed with or without bootstrapping in case of Euclidean trees. We have also constructed n-dimensional tree based on a new algorithm (see methods) to construct trees based on N dimensions (Milner Kumar & Sohan P. Modak, manuscript in review). With the exception of the position of *Leptospira* (Spirochaetes) and *Acinetobacter* (Proteobacteria), all tree topologies were identical. In case of bootstrapped Euclidean trees *Leptospira interrogans* appeared near the base of the tree whereas Acinetobacter forms a clade with Betaproteobacteria. In trees without bootstrapping, *Leptospira* and *Acinetobacter* interchange their positions with *Leptospira* being at the root of Epsilongproteobacteria (Fig. 3; S4 Figure).

## Comparison of topologies of single or multiparametric trees with 16S rDNA tree and estimation of Taxonomic fidelity (F) and Robinson-Foulds metric (RF)

We have used a novel taxonomic fidelity algorithm (see methods) to estimate the extent of similarity between the topology of benchmark taxonomic tree built for 119 species and that of phylogenetic trees for individual tRNA synthetases, consensus Euclidean tree and those using concatenated sequences. The Fidelity algorithm carries out clade by clade comparison among two tree topologies. We find that the Fidelity value (F) falls in the range 83.5% to 92.9% for 15 individual tRNA synthetase trees (Table 1). In contrast, the ‘F’ values for concatenated trees are within the range, 88.9% to 89.6%, while for all trees based on original or bootstrapped Euclidean distances generated is even superior at 90.3%. The ‘F’ values for all taxonomic groups and for all trees are provided as S1 Table. However, the 16S rDNA tree gives a considerably lower ‘F’ value of 87.2% (Table 1). Furthermore, in this tree, *Picrophilus torridus* (superkingdom Archaea) did not group with other species from Archaea (Fig. 4; S5 Figure). It is notable that species under the Phylum Cyanobacteria do not form a single clade for 16S rDNA based tree. At the genus level, all species under genera *Vibrio*, *Salmonella*, *Pseudomonas* and *Prochlorococcus* did not form independent unique clades. Yet all species in the genus *Bacillus* formed discrete clade. Members of phylum Spirochaetes (*Borrelia burgdorferi* and *Leptospira interrogans*) also formed a unique clade (Fig. 4; S5 Figure). It is notable that 33 taxa from the phylum Firmicutes exhibit dispersed topology and variable taxonomic fidelity. For example, 8 species from the order Lactobacillales and 16 from the order Bacillales show poor ‘F’ value 62.5% and 83.3%, respectively (S1 Table). *Wolinella succinogenes* was always found to intersect the *Helicobacter* genus clade in all multidimensional and most of the single gene trees, but in the case of 16S rDNA tree, *Helicobacter* formed a single clade.

**Figure 4:**
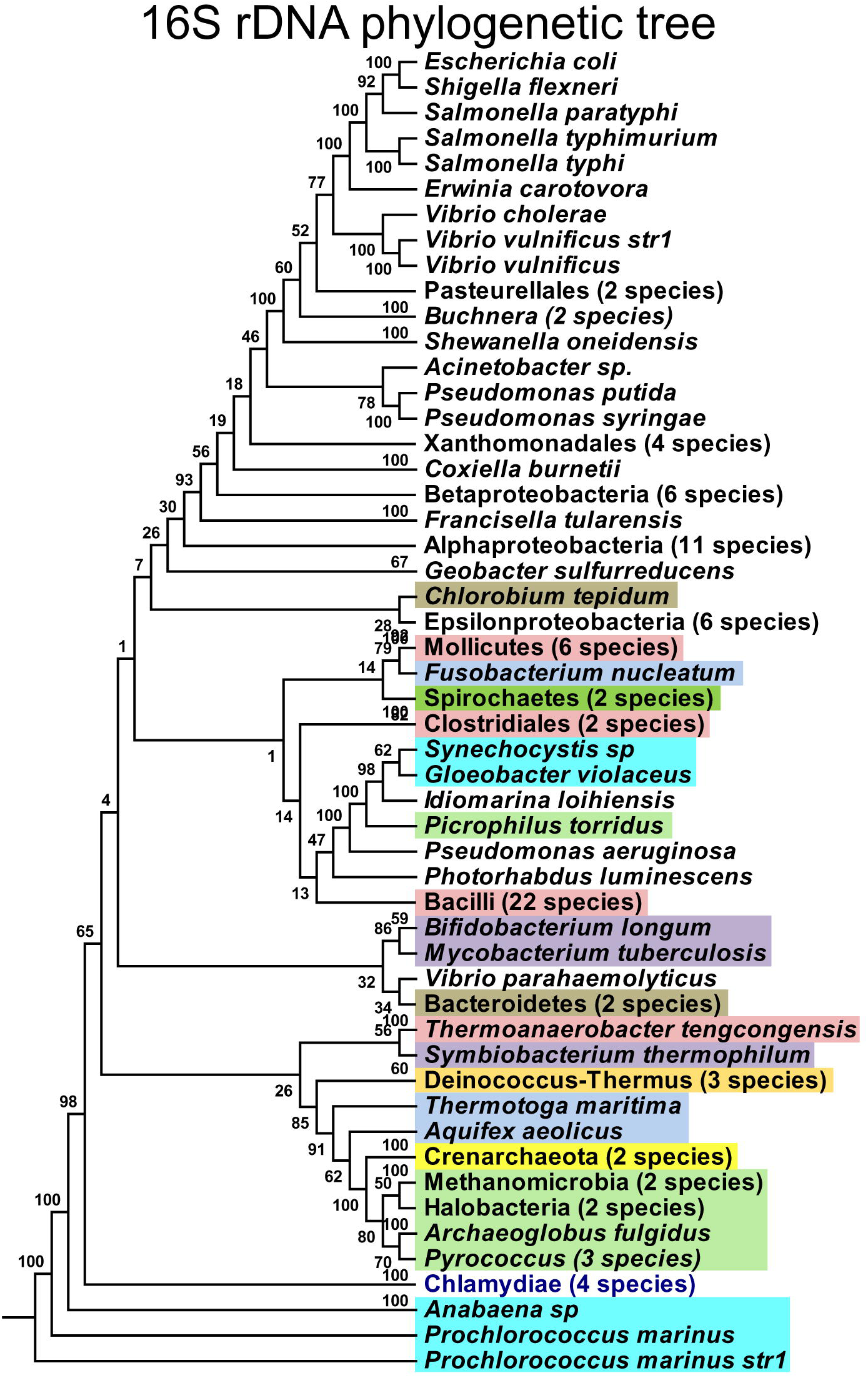
Phylogenetic tree for 16S rDNA sequences. rDNA sequence data for 116 out of 119 available species was retrieved and phylogenetic trees were constructed as described in methods section. The expanded version of tree is provided as Supplementary Figure 5.

We also estimated the Robinson-Foulds metric (RF) [16] between phylogenetic trees and the taxonomic tree and compared these to ‘F’ values (Table 1). We find these two metric are inversely proportional.

## Discussion

Constructing phylogenetic trees by comparing the nucleotide or amino acid sequence of a given gene/polypeptide among a set of species is useful in determining their relative evolutionary positions. However, it is widely noticed [3,13] that for the same set of species the topology of phylogenetic trees varies from one gene/polypeptide to another. This is understandable since the mutation rates differ from one gene to another [14]. Indeed, due to the multi-genic composition of each organism, the most reasonable phylogenetic tree should take into account a consensus of representative genes or phenes, which would be concordant with the original phylogenetic tree structure described by Darwin, based on morphological traits that are necessarily polygenic. Hence, there is a need to construct phylogenetic trees based on multiple traits that offer a consensus solution. It is also important to select appropriate genes for such analysis. We selected aminoacyl-tRNA synthetase for the current study, since these genes are involved not only in analogous function but probably in one of the most ancient [17] and pivotal steps to recruit specific amino acids for transport to the site of translation.

A relatively recent method to examine phylogeny using multiple genes involves using concatenated gene sequences. However, we felt that the manner in which many molecular sequences are concatenated may be as important as their evolutionary relevance. For example, even for a single gene or phene the nucleotide/amino acid sequences differ in chain-length among different species which could create problems in their alignment. Therefore, in the concatenation method as described routinely, sequences are aligned first and then concatenated. We have tackled this problem by applying MSA either before or after concatenation, by concatenating in alphabetical order, or by concatenating in the order of chain length.

Unlike the use of the term concatenation for megapolynucleotide/polypeptides created by end-to-end joining, concatenation here means formation of a chain link, as in the case of chain links of multiple supercoiled DNA molecules. For example, megapolypeptides were generated *in silico* by joining the carboxy terminal amino acid of the first polypeptide to the amino terminal of the second polypeptide, then second to third and so on. The same method will apply for 3’ - 5’ joining of polynucleotides. When for the same polypeptide the sequence length varies among different species, its joining to the sequence of the next polypeptide may lead to misalignment, a process that will amplify with increasing numbers, types of polypeptides, and species and causes loss of information in the final tree in addition to the alignment process becoming computationally expensive. It is known that different MSA protocols do exhibit varying accuracy and time complexity [15].

When we ligated (concatenated) the amino acid sequences of 15 aminoacyl-tRNA synthetases in the increasing order of chain lengths, followed it by the sequence alignment of these megapolypeptides, the resulting tree topology was comparable to that of concatenates formed after individually aligning polypeptide sequences first (Fig. 2B and 2D). Among the three types of protocols for concatenation prior to MSA, the tree topologies indeed differ in the position of Gamma-and Beta-proteobacteria (Fig. 2A-C). In any case, the phylogenetic trees built for these 3 types of concatenates exhibit comparable taxonomic fidelity values.

When comparing the extent of sequence homology for a given gene or polypeptide, one should recognize that across a set of species these would bear some sequence segments that are highly conserved (Block) and others that vary (non-Block). When we first aligned aminoacyl-tRNA synthetases individually and then concatenated these, the tree topology of non-conserved (non-Block) sequence concatenates is similar to that for total sequences and considerably different from the tree for conserved (Block) sequence concatenates (S4 FigureS4 Figure). This result is surprising because one would have expected trees for total sequences to exhibit less variability in topology than non-blocks. We suggest that it is the variable (non-block) sequence that is relevant to the extent of closeness in phylogenetic analyses.

In 2003, Milner et.al. [9] discovered a novel way of generating all-pairs distance coordinates simultaneously for 3 types of traits by using Euclidean geometry wherein the extent of similarity among a set of species for one trait is plotted on x-axis, second on the y-axis and third on z-axis. This process generates a graphical representation of the extent of similarity among the pairs of species/objects in a 3D space. In that study, they examined the phylogenetic relationships based on immuno-crossreactivity of lens crystallins from 8 species against three different anti-crystallin antisera. Subsequently [9,10], in addition to the immuno-crossreactivity, they also used RNA and DNA nucleotide sequences to estimate all pairs Euclidean distances to construct phylogenetic trees. In their studies [10] even immuno-crossreactivity of polyclonal antibodies to lens crystallins from three mammalian species revealed that some of the chiropteran species classified as microchiroptera should actually be classified among megachiropterans, which was also consistent with their feeding habits and lack of echo-location apparatus.

This method was further refined to compare the phylogenetic tree topologies of three mitochondrial polypeptides from 76 species and it was found that the taxonomic fidelity substantially improved in the consensus tree constructed using Euclidean geometry [17]. Furthermore, contrary to the claims made on the basis of phylogenetic trees using concatenated sequences of nuclear transcripts by Delsuc [2], hemichordates and cephalochordates along with echinoderms appear at the root of the phylum chordata, i.e. just before cyclostomes, while urochordates occupy a considerably lower position along with proteostomes [11].

Here, we have used this method to generate consensus phylogenetic trees step-wise using amino acid sequences of 15 tRNA synthetase polypeptides from 119 prokaryotes. It should be noted that the all pairs distances found in each set of 3 polypeptides are then used along with two similar sets for different polypeptides along x-, y-and z-axes to obtain a distance matrix first for 9 proteins and eventually for all 15 polypeptides from the same 119 prokaryotes. We find that tRNA synthetases - a discrete set of proteins involved in one of the most crucial steps in transfer of genetic information from gene to phene, show considerable phylogenetic diversity among prokaryotes. The phylogenetic tree using the Euclidian geometry offers a consensus topology, which is independent of their chemical or physical properties and the order in which these proteins are grouped.

From Euclidean trees, a surprising observation concerns the topological position of the class Alphaproteobacteria that form a clade with Archaea, rather than the remaining Proteobacteria (Fig. 3). Even in all types of trees for concatenated sequences, Alphaproteobacteria form a separate clade dissociated from the rest of the species in the phylum Proteobacteria and indeed closer to Archaea and Firmicutes. (Fig. 2) In contrast, in the 16S rDNA phylogenetic tree and NCBI taxonomic trees, most species from the phylum Proteobacteria form a clade (Fig. 4; S6 Figure). Thus, based on our multidimensional phylogenetic trees, the taxonomic position of class Proteobacteria needs to be revisited.

While it is possible to estimate distance between any two trees [16], these do not allow validation of phylogenetic tree topologies against the taxonomic hierarchy of the species under investigation. We have used different methods to construct a large number of phylogenetic trees and it is difficult to validate the composition of clades for species under similar taxonomic clusters. In any case, the reasonableness of any phylogenetic tree should be judged on the basis of its relative concurrence with the taxonomic tree as a benchmark for a corresponding set of taxa. Therefore, we have used the taxonomic fidelity algorithm developed by Milner Kumar & S. P. Modak, (manuscript in review), which quantitatively examines all clades for clade-by-clade similarity and positional correspondence between a given phylogenetic tree and the taxonomic tree for same set of species. We find that phylogenetic trees for individual aminoacyl-tRNA synthetases exhibit ‘F’ values ranging from 83.5% to 92.9%, while both 3 and 15-dimensional Euclidean phylogenetic trees (Fig. 3) offer a stable fidelity value (90.3%). Phylogenetic trees constructed from tRNA synthetases concatenated using different methods also provide comparable fidelity (F) values (88.9% to 89.6%) (Table 1, S1 Table). In addition, tree topologies for Euclidean distances are not only identical to those even after bootstrapping but also independent of the method of grouping the polypeptides. This is unlike the situation where tree topologies change with the order in which sequences are concatenated (Fig. 2 and 3) and suggests that Euclidean distances actually offer a better approximation of all-pairs phylogenetic distances. Another argument favoring construction of multiparametric phylogenetic trees using Euclidean geometry stems from the fact that significant differences appear in tree topologies of concatenated trees for ‘Block’ and ‘Nonblock’ sequences.

Improving upon the earlier studies [9–11], we have now extended the phylogenetic tree constructs to represent consensus of 15 proteins. Furthermore, we have applied yet another algorithm to construct trees based on N dimensions/parameters (Milner Kumar & S. P. Modak, Manuscript in review).The resultant tree, obtained in a single step, is identical with those using serial groupings of Euclidean distances for 3 traits (Fig. 3). Clearly, the method can be further extended to include sequence data for a consortium of large numbers of genes/polypeptides to obtain a polygenic/polyphenic consensus phylogenetic tree. One of the striking examples of improved tree topology using Euclidean geometry involves construction of the tree for Asp, Cys and Ser tRNA synthetases (group A based on the mean of highest distance), which shows that as compared to the single gene trees, exhibiting ‘F’ values of 87.3%, 84% and 89.8%, respectively, taxonomic fidelity of multidimensional tree was 90% (S1 Table).

Robinson-Fould’ s ‘RF’ metric [16] compares topology of any two trees and estimates the extent of dissimilarities while, in our modified algorithm, ‘F’ value estimates the closeness of a ‘test’ tree to the benchmark taxonomic trees. ‘RF’ compares two trees and the score is based only on identical clades. In contrast, our taxonomy fidelity value ‘F’ seeks best approximation between every clade in the Test tree and the corresponding one in the benchmark tree and assigns at least a fractional value. Thus, in the worst case scenario, ‘F’ captures even the minimal similarity between a clade-pair, which ‘RF’ does not. We find that the values of ‘F’ and ‘RF’ are inversely proportional. In this paper, we have estimated both ‘RF’ and ‘F’ and we find that ‘F’ not only assesses the reasonableness of consensus tree against taxonomic trees, but further offers higher resolution than ‘RF’.

According to Woese et al [17] half of the aminoacyl-tRNA synthetases agree with established classical phylogeny while, the rest contradict it, although, most of major taxonomical groupings are maintained in all aminoacyl-tRNA synthetases. In our studies reported here, minor differences at taxonomic levels in few or more single gene trees were taken care of in case of multidimensional trees as depicted in supplementary table, where several taxonomic definitions show variable ‘F’ values for single gene trees but in case of multidimensional trees these form a single clade. Classes such as Thermoprotei, Bacilli and Clostridia show variable clustering patterns in single gene trees, but form a single clade in case of multiparametric trees.

Excepting Leu-tRNA synthetase, in all single gene trees, *Borrelia burgdorferi* and *Leptospira interrogans* (phylum Spirochaetes) form a separate clade. This is in conflict with the 16S rDNA tree in which they form single clade.

Although a large number of 16S rDNA sequences are available and used to fine-tune microbial taxonomy [18], using these to identify otherwise unknown species under the genus Bacillus have been only partially successful [19] and obviously need to be complemented with many protein coding gene sequences that can act as distinguishing traits, thereby leading to a consortium of multiple molecular identifiers. Clearly, the 16S rDNA tree cannot be routinely used as a benchmark. In fact, it is clear that the fidelity of 16s rDNA tree is considerably inferior to that of multiparametric trees based on either Euclidean geometry or concatenation.

According to NCBI’ s latest classification [20], the genera *Mycoplasma*, *Mesoplasma* and *Ureaplasma* are grouped under phylum Tenericutes and separate from Firmicutes (S6 Figure), which is confirmed by the topologies of our tree for Euclidean distances as well as end-to-end joined sequences (Fig. 2 and 3). Furthermore, in new NCBI classification [20], *Symbiobacterium thermophilum* is placed in the order Clostridiales, phylum Firmicutes (S6 Figure), rather than under phylum Actinobacteria [21]. We also find that *Symbiobacterium thermophilum* is positioned away from the rest of the Actinobacteria and clusters with Firmicutes (Fig. 3). We conclude that the Euclidean distance trees based on multiple traits such as amino acid sequences of 15 tRNA synthetases offer a consensus hierarchy that closely resembles the new NCBI taxonomy based on polygenic traits and questions the logic of reliance on using rRNA as indicators for microbial taxonomy.

During the evolution of the living systems after the establishment and consolidation of the molecular hierarchy between DNA as the carrier of genetic information and polypeptides as the final structural and functional phenotypes, a multi-step process and a battery of molecules are required to ensure the transliteration from genes to phenes. Clearly the genetic code must have co-evolved with the transfer RNAs and tRNA synthetases required to link these to corresponding amino acids. Thus, tRNA synthetases have probably evolved earlier than ribosomal RNA.

Aminoacyl-tRNA synthetases represent universal, constant and ancient functions defined mainly by the tRNAs that are among the most ancient and structurally constant molecules in the cell [17]. Obviously, synthetases collectively carry a record of far earlier times that preceded those represented by the root of the universal (rDNA) phylogenetic tree [17].

From the sequence-based analysis, events that gave rise to Leu-, Ile-, and Val-tRNA synthetase [22] and Trp-and Tyr-tRNA synthetase [23] cluster monophyletically with respect to amino acid specificity. In contrast, from the topologies of single gene trees, we find that Taxonomic Fidelity value, ‘F’ is highest for Leu-tRNA synthetase and lowest for Val-tRNA synthetase with the remaining 3 molecules holding intermediate positions (Table 1). In the Euclidean distance tree based on 15 Aminoacyl-tRNA synthetases the consensus F value is 90.3% which is only lower than that for individual Leu-and Lys-tRNA synthetases.

Single gene phylogenetic trees, including those for rDNA, constructed by comparing a single trait and cannot be expected to mimic taxonomic trees. The taxonomic relationships are based on several morphological and physiological characteristics that are consequent to the expression of large number of genes. In the present study, we have used 15 aminoacyl-tRNA synthetases to construct multiparametric phylogenetic trees and find that the phylogenetic hierarchy based on tree topology from Euclidean geometry for 119 prokaryotes is closer to the taxonomic hierarchy than that for individual proteins.

## Conclusions

We have constructed multi-genic trees based on Euclidean distances among molecular sequence alignments. For those constructs, we aligned sequences of 15 aminoacyl-tRNA synthetases from 119 species and constructed a tree consensus for the 15 phenes/polypeptides. We used an algorithm that allows construction of a phylogenetic tree based on N-dimensions wherein each dimension is ascribed to sequence alignment of a given polypeptide. Using this algorithm, we have constructed an N-dimensional phylogenetic tree of 15 tRNA synthetase sequences from 119 prokaryotes. Comparison of the taxonomic fidelity shows that ‘F’ values are 87.2% for 16s rRNA, range between 83.5% to 92.9%, for individual aminoacyl-tRNA synthetases, are 89% for trees based on concatenates of 15 polypeptide sequences and are even higher (90.3%) for those based on N-dimensional tree topology. Thus, N-dimensional Euclidean tree gives the best consensual topology.

From Euclidean trees, a surprising observation reveals that the class Alphaproteobacteria forms a clade with Archaea, rather than the remaining Proteobacteria. Even in trees for concatenated sequences, Alphaproteobacteria form a separate clade dissociated from the rest of the species in the phylum Proteobacteria and indeed closer to Archaea and Firmicutes. In contrast, in 16S rDNA phylogenetic tree and NCBI taxonomic trees, most species from the phylum Proteobacteria form a unique clade. Thus, based on our multidimensional phylogenetic trees, the taxonomic position of class Proteobacteria needs to be revisited.

## Materials and methods

### Data collection

Aminoacyl-tRNA synthetase sequences were downloaded from PIR, PRF, Swissprot and Tremble (www.genome.jp) databases using in-house python scripts. From the protein databases, 5948 polypeptide sequences were retrieved representing 505 species and 20 aminoacyl-tRNA synthetases including 18 monomeric polypepetides (Ala, Arg, Asn, Asp, Cys, Glu, Gln, His, Isl, Leu, Lys, Met, Pro, Ser, Thr, Try, Tyr and Val) and, dimeric synthetases Phe and Gly, each with α and β chains. We filtered out pseudo, hypothetical and predicted protein sequences and found that amino acid sequences of fifteen aminoacyl-tRNA synthetases, viz; Arg, Asp,Cys, Isl, Leu, Lys, Met, α chain Phe, β chain Phe, Pro, Ser, Thr, Try, Tyr and Val were available in only 119 species (listed in S2 Table). The data used here ensures optimal representation of maximum number of taxonomic groups with given polypeptide sequences. Out of these 119 species, we found 16S rDNA sequences for only 116 species and these were retrieved. The taxonomic classification and the tree for above 119 species was retrieved from NCBI [20,24]. Fig. 5 depicts the methods used to generate single or multi-gene phylogenetic trees.

**Figure 5:**
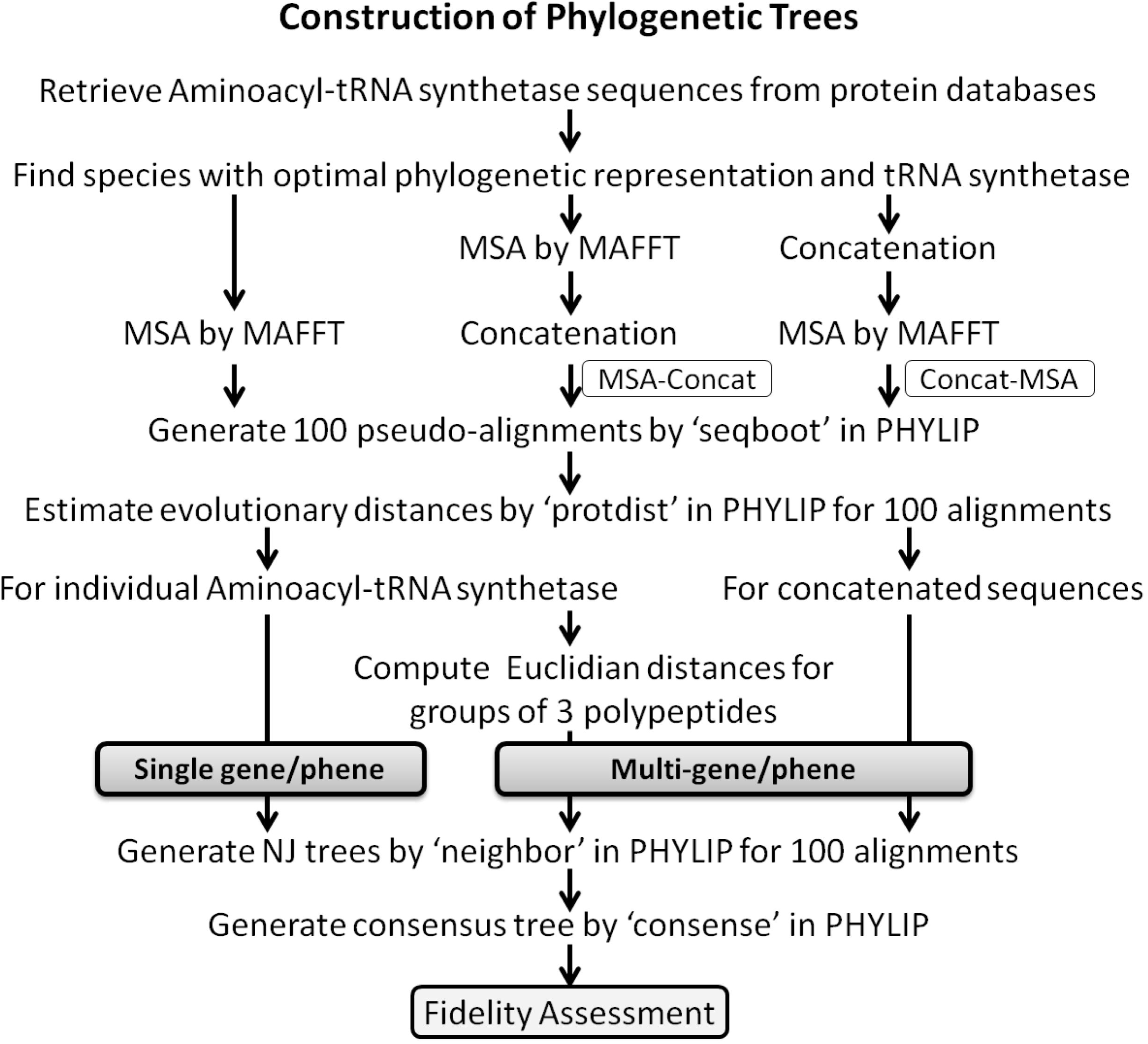
Construction of single or mulit-gene phylogenetic trees. The figure describes overall steps carried out in this study to generate single or multi-gene trees.

### Construction of phylogenetic trees

The sequences were aligned using MAFFT [25] and ClustalW [26]. For each polypeptide from 119 species, we used the program “seqboot” in PHYLIP [27] to generate 100 pseudo alignments from which distance matrices were computed using “protdist” in PHYLIP [27]. Neighbor-joining trees [28] were then computed using the program “neighbor” [27] followed by the program “consense” in PHYLIP [27] to construct the bootstrapped consensus tree [29].

### Consensus phylogenetic trees using concatenated sequences

For concatenation, the sequences were end-end joined between the carboxy terminal of the first polypeptide and the amino terminal of the second polypeptide, then second to third and so on till a 15 polypeptide long megapolypeptide was obtained for each species. Aminoacyl-tRNA synthetase sequences were concatenated using four different protocols: (1) concatenation in their alphabetical order (alphabetical order concat **→**followed by MSA); (2) concatenation in the order based on increasing average chain length **→**followed by MSA; (3) concatenation in the order of decreasing average chain length **→**followed by MSA) (Table 2); (4) MSA of individual sequences from all species first followed by the end-to-end joining (concatenation) of the aligned sequences in an alphabetical order (MSA **→**concat) to overcome the effect of sequence length variations. The phylogenetic trees were then computed as before.

**Table 2:**
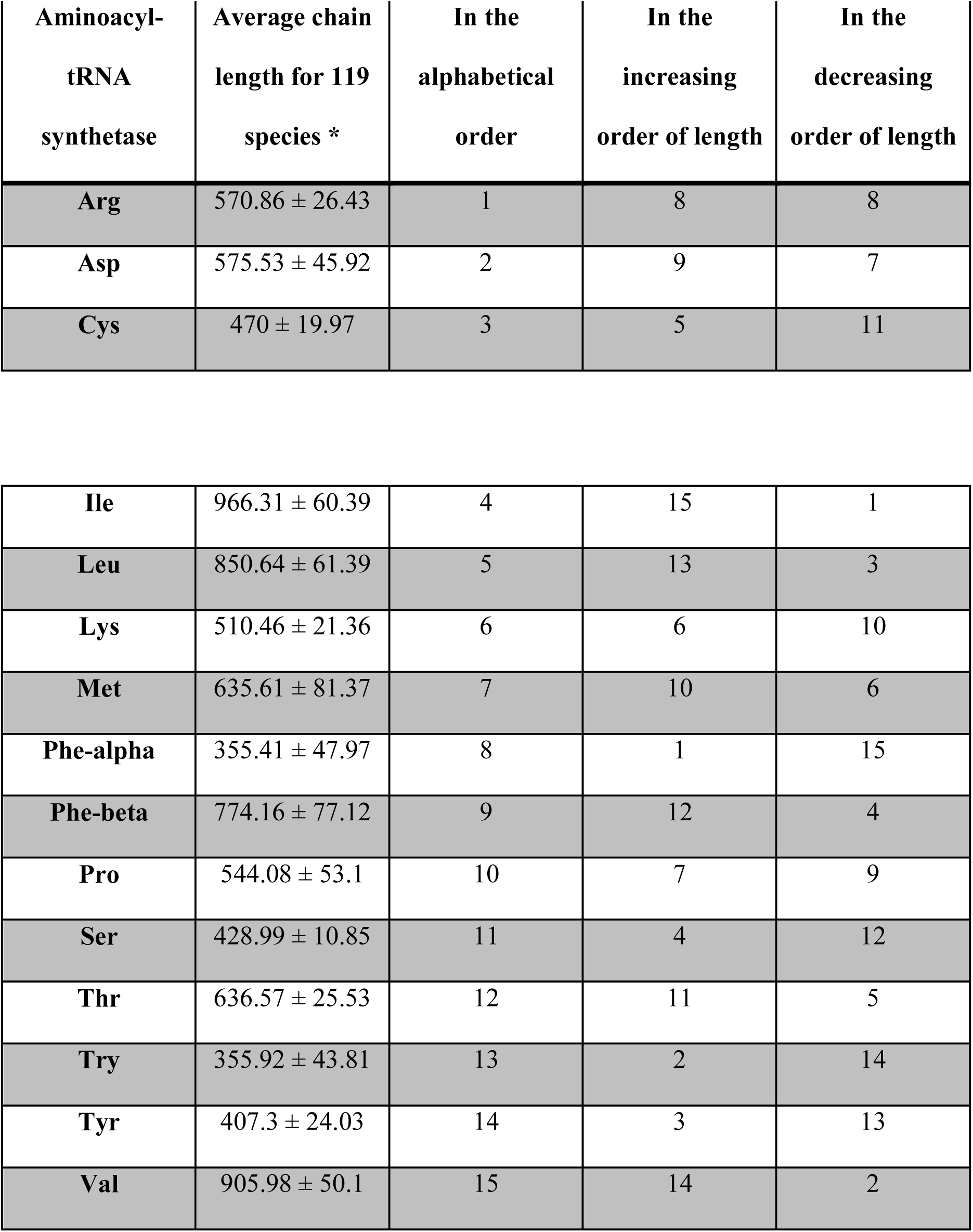
The serial order for concatenation of aminoacyl-tRNA synthetase sequences. Standard deviation for chain length (number of amino acid residues) among 119 species.

In another set, all four types of the concatenated megapolypeptides obtained above were subjected to the program “GBlocks” [12] and the output of contiguous stretches of conserved (Blocks) and non-conserved (non-Blocks) sequences were separated and used to construct phylogenetic trees as before.

### Consensus phylogenetic trees based on Euclidean distances

From the MSA for individual tRNA synthetase sequences from 119 species, the aligned sequences were subjected to PROTDIST and the pair-wise distances for three synthetases were estimated using All Pairs Shortest Distance Algorithm [3] and plotted the on x-, y-and z-axis, respectively. In each plot, a species was placed at the origin (0, 0, 0) and with the known pair-wise distances and the remaining 118 species were positioned at their corresponding positions along x, y and z axes in the 3D space [9–11]. This process was repeated by sequentially replacing the species at the origin in turn to eventually generate 119 3-D plots. From these plots we estimated all-pairs Euclidian distances between the species at the origin and those in the 3-D space [9,10] which were then used to construct phylogenetic trees.

The method to estimate all-pairs Euclidean distances in a 3D space uses only 3 traits/characters/molecular types. Therefore, the procedure was reiterated by creating 5 groups of 3 tRNA synthetase polypeptides (Fig. 6) based according to the chemical properties of the acceptor amino acids, *viz* charged amino acids (Arg, Asp and Lys), polar uncharged (Pro, Ser and Thr), non-polar aliphatic (Ile, Leu and Val), non-polar aromatic (α chain Phe, β chain Phe and Tyr) and mixed (Cys, Met and Try) (Fig. 6A). The remaining two groups were based on the extent of conservation in terms of all pairs distances among aminoacyl-tRNA synthetase for 119 species as determined from the distance matrices obtained using TREE-PUZZLE [30] for each individual aminoacyl-tRNA synthetase. The all-pairs distances were used as measure for the extent of conservation and were used in the remaining two groupings, as with (b) increasing order of sum of all pairs distance for each aminoacyl-tRNA synthetase (table 3, Fig. 6B), and (c) the mean of highest all pairs distance for each species for individual aminoacyl-tRNA synthetase was estimated and the proteins were grouped in their increasing order (Table 3, Fig.6C).

**Figure 6:**
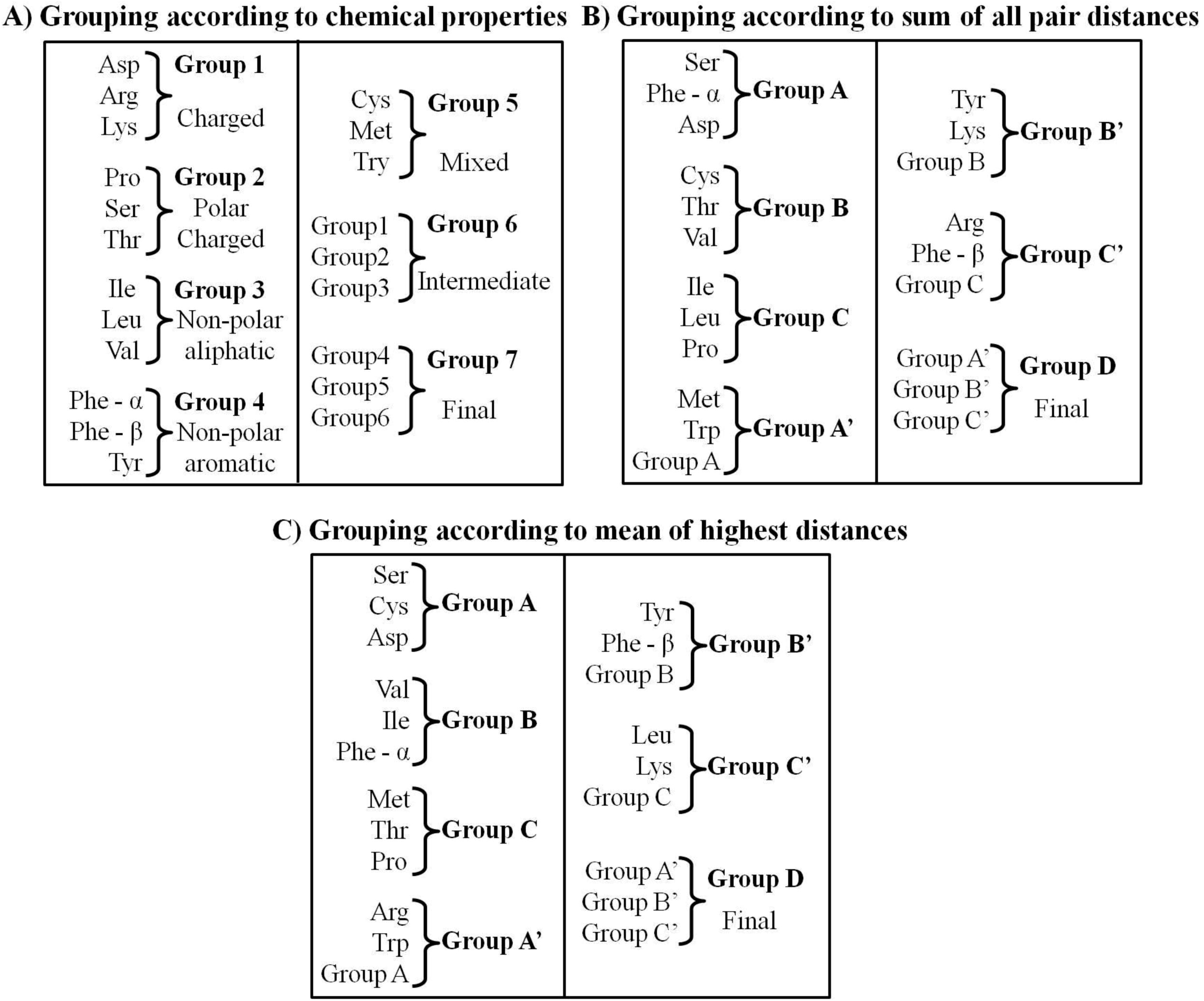
Grouping strategies used for constructing multidimensional trees in 3D space. A) Aminoacyl-tRNA synthetases were divided in five groups based on chemical properties. Distance matrices generated from these groups were further grouped to generate intermediate and final trees. B) The grouping was performed based on sum of evolutionary distances as described in methods. C) Mean of the highest all pairs distances were used for classifying the aminoacyl-tRNA synthetases.

**Table 3:** Evolutionary distances estimated by TREE-PUZZLE

| **Aminoacyl-tRNA**<br>**synthetase** | **Sum of distances** | **Mean of highest distance** |
| --- | --- | --- |
| **Arg** | 21026 | 2.164 |
| **Asp** | 12532 | 1.523 |
| **Cys** | 13577 | 1.48 |
| **Ile** | 16387 | 1.772 |
| **Leu** | 16509 | 2.466 |
| **Lys** | 20900 | 3.834 |
| **Met** | 17862 | 1.838 |
| **Phe-alpha** | 13450 | 1.775 |
| **Phe-beta** | 20906 | 2.411 |
| **Pro** | 17124 | 2.146 |
| **Ser** | 12077 | 1.332 |
| **Thr** | 13889 | 1.983 |
| **Try** | 17257 | 2.317 |
| **Tyr** | 18850 | 2.343 |
| **Val** | 14303 | 1.719 |

From the Euclidean distance matrix all pairs Euclidean distances for each of the 5 groups were merged sequentially (Fig. 6) to construct a consensus tree for 15 tRNA synthetases. From aligned sequences for individual types of aminoacyl-tRNA synthetase of 119 species, we also generated 100 pseudo alignments using the program “seqboot” [27] and the corresponding 100 distance matrices were estimated using “protdist” [27]. These were then merged as before to obtain a unique 2-D multidimensional tree representing a bootstrapped Consensus for 15 Aminoacyl-tRNA synthetases in 119 species [29].

### N-Dimensional consensus Euclidean phylogenetic tree

15-dimensional Euclidian distance matrix can be estimated from 15 evolutionary distance matrices using the following equation (Milner Kumar & Sohan P. Modak, manuscript in review):

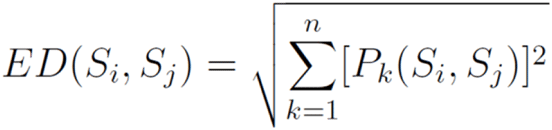

Where, ED(S_i_, S_j_) - 15-dimensional Euclidian distance between species S_i_ and S_j_. The values of i, j vary in between 1 and the number of species used in the study. P_k_(S_i_, S_j_)-distance between species S_i_ and S_j_ from P_k_ i.e., the k^th^ parameter/dimension/phene.

### Taxonomic fidelity

Using the clustering algorithm for taxonomic fidelity (Milner Kumar & Sohan P. Modak,(manuscript in review), we have compared the extent of clade-clade correspondence in the topology of taxonomic tree for 119 species, derived from the NCBI taxonomy browser [20,24] and each single gene and multidimensional tree (Fig. 5). In this comparison, we used only those taxonomic groups represented by at least two species (S2 Table). The taxonomic fidelity “F” is estimated using the equation:

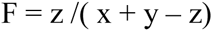

where, F–the fidelity, z–number of species common to the taxonomic clade/s and the corresponding phylogenetic clade/s, x–number of species in “Taxonomic Clade”, y– number of species in “Phylogenetic Clade”. Taxonomic fidelity ‘F’ was estimated for each taxonomic group. Sum of F value (depicted as per cent of total number of taxonomic groups) for each tree indicates its similarity or closeness to the classical taxonomic tree. With F equal to 1, the clade mimics the taxonomic cluster; with F less than 1, the cluster may either be incomplete due to absence of related species, or has acquired a taxonomically unrelated species, or both.

### Robinson-Foulds metric

We have also estimated the extent of dissimilarities ‘RF’ between the phylogenetic trees and benchmark taxonomic tree by Robinson-Foulds method [16] for 119 species using the equation

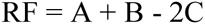

Where, ‘A’ is the total number of clades in first tree, ‘B’ is the total number of clades in second tree and ‘C’ is the number of clades common to both trees. When ‘RF’ equal to 0, then both the trees are identical. If two trees do not share any common clade/s then ‘RF’ will be the sum of all clades in both trees. This is the maximum value ‘RF’ can attain. In this method, ‘RF’ values are bi-directionally identical.

## Acknowledgements

Authors would to like to acknowledge Swanand Joshi, Paresh Vishwasrao and Jai Kulkarni for help in data mining and critical discussions. We are grateful to Glynnis Kirchmeier for a critical review of the manuscript.

---

Supplementary Fig. 1: **Single gene phylogenetic trees for aminoacyl-tRNA synthetases**. The figure depicts expanded version of the single gene phylogenetic trees for 15 polypeptides.

Supplementary Fig. 2: **Phylogenetic trees for concatenated aminoacyl-tRNA synthetases sequences**. The figure depicts expanded version of the concatenated phylogenetic trees.

Supplementary Fig. 3: **Phylogenetic trees for Blocks and Non-blocks regions of concatenated aminoacyl-tRNA synthetases sequences**. Block and Non-block regions were retrieved after the alignment of concatenated sequences, and phylogenetic trees were constructed as described in methods section.

Supplementary Fig. 4: **Multidimensional phylogenetic trees based on Euclidean distances**. Trees were constructed with (A) and without (B) bootstrapping. The value on the branch indicates the frequency of occurrence from 100 bootstraps.

Supplementary Fig. 5: **Phylogenetic tree for 16S rDNA sequences**. The expanded version of the phylogenetic tree based on 16S rDNA sequences.

Supplementary Fig. 6: **Taxonomic phylogenetic tree using NCBI**. All 119 species were submitted as input for the NCBI taxonomy server and a tree was generated for these species.

